# Glutathione S-Transferase Genotype polymorphism and Primary Open-Angle Glaucoma in an Italian population

**DOI:** 10.1101/413435

**Authors:** Sergio C Saccà, Carlo A Cutolo, Stefano Gandolfi, Giorgio Marchini, Luciano Quaranta, Alberto Izzotti, Tommaso Rossi, Carlo Enrico Traverso

## Abstract

**Purpose:** Oxidative damage to the trabecular meshwork (TM) represents one of the pathogenic mechanisms leading to primary open angle glaucoma (POAG). Glutathione S-transferase mu 1 (GSTM1) may neutralizes reactive oxygen species protecting the TM. The present paper investigates the prevalence of GSTM1 null genotype in an Italian population, and its association with POAG treated either medically or surgically.

**Methods:** In a case-control study, the GSTM1 genotype was identified in POAGs and controls. The POAGs patients were divided in two groups: medical POAGs and surgical POAGS. Medical POAGs consisted of patients with a well-controlled intraocular pressure (IOP) by IOP-lowering medications and a stable visual field (VF). Patients with an uncontrolled IOP and a progressing VF that were submitted to incisional surgery formed the surgical POAGs’ group.

**Results:** We enrolled 104 medical POAGs, 158 surgical POAGs and 263 Controls. No significative differences between the groups existed regarding age and gender (p=0.275 and p=0.950, respectively). All the enrolled subjects were Caucasian of Italian descents. The GSTM1 null genotype was identified in 57 (45.2%) medical POAGs, 91 (57.6%) surgical POAGs and, 119 (45.3%) controls (p=0.033). The association between medical POAG and GSTM1 null status was non-significant (OR= 1.44, 95% IC = 0.86 to 2.39) whereas the association was significant for surgical POAGs (OR= 2.01, 95% IC= 1.26 to 3.21)

**Conclusions:** Our results showed an association between the GSTM1 null genotype and glaucoma that require surgery in an Italian population. GSTM1 null genotype detection may help to identify high-risk glaucoma patients that require a closer follow-up and a more aggressive treatments.

## Introduction

According to the Free Radical Theory of Aging, aging and its related diseases are the consequence of damage by Reactive Oxygen Species induced in cell structures and the inability of endogenous antioxidant defenses to offset these changes. ROS attack on macromolecules is often called oxidative stress. The ROS produced during aerobic respiration have deleterious effects on cell components and connective tissues, causing cumulative damage over time that eventually results in aging and death.[1] Most of the endogenous free radicals are produced by mitochondria, and most of the free radical damage is to mitochondrial membranes and mitochondrial DNA. Mitochondria generate a significant amount of cellular energy through the consumption of most of the intracellular oxygen. Indeed, about 90% of cellular oxygen is consumed within the mitochondria, particularly in the inner membrane, where oxidative phosphorylation occurs[2]. Oxidative damage induces mitochondrial stress, which over time, becomes damage: damaged mitochondria become progressively less efficient, losing their functional integrity and releasing more oxygen molecules, thereby increasing the oxidative damage and finally leading to the accumulation of dysfunctional mitochondria severe levels of mitochondrial damage trigger apoptosis through the intrinsic activation pathway[3].

In the eye, as in several neurodegenerative diseases, the normal antioxidant defense mechanisms decline, which increases the vulnerability of the brain and its eversion or eye to the deleterious effects of oxidative damage[4]. It is thought that free radicals of mitochondrial origin are among the leading causes of mitochondrial DNA damage (mtDNA). Indeed, high levels of 8-hydroxy-2’- deoxyguanosine (8-OHdG), a biomarker of oxidative damage to DNA, in the mtDNA of the aged brain and, in the trabecular meshwork (TM) occur[5,6]. In glaucoma (POAG and SPEX), the damage to the trabecular mtDNA is particularly severe in comparison with controls, and results in a striking increase in mtDNA 4977 deletion[7]. Primary open-angle glaucoma (POAG) is a multifactorial disease that ultimately leads to apoptosis of retinal ganglion cells (RGC); oxidative damage plays an important role in its pathogenesis, as also do genetic factors[8,9]. Indeed, several studies have indicated that some cases of glaucoma are hereditary, despite the fact that no single-hit POAG gene has been identified, multiple genetic risk factors probably contribute to its pathogenesis. Genes known to contribute to glaucoma are optineurin[10], myocillin[11], and TANK-binding kinase 1 (TBK1)[10], furthermore, recent genome-wide association studies have identified several single nucleotide polymorphisms to be associated with POAG (Table 1). In the general population, these gene mutations are approximately >5–10% of all POAG cases[12]. A genetically determined phenomenon that may be involved in the pathogenesis of POAG, which could have a higher impact, is the lack of glutathione transferase isoenzyme M1 (GSTM1). Indeed, several studies have indicated the importance or otherwise of the GSTM1 null genotype, depending on the patient populations studied[13]. GSTM1 catalyzes the detoxification of electrophilic reactive oxygen species by conjugation with reduced glutathione. Two polymorphic variants occur, i.e. normal-function (wild type) or loss-of-function due to homozygous deletion on both alleles (null genotype). The existence of a single heterozygous deletion does not result in a loss of function and is thus regarded as wild-type normal polymorphism.

**Table 1:**
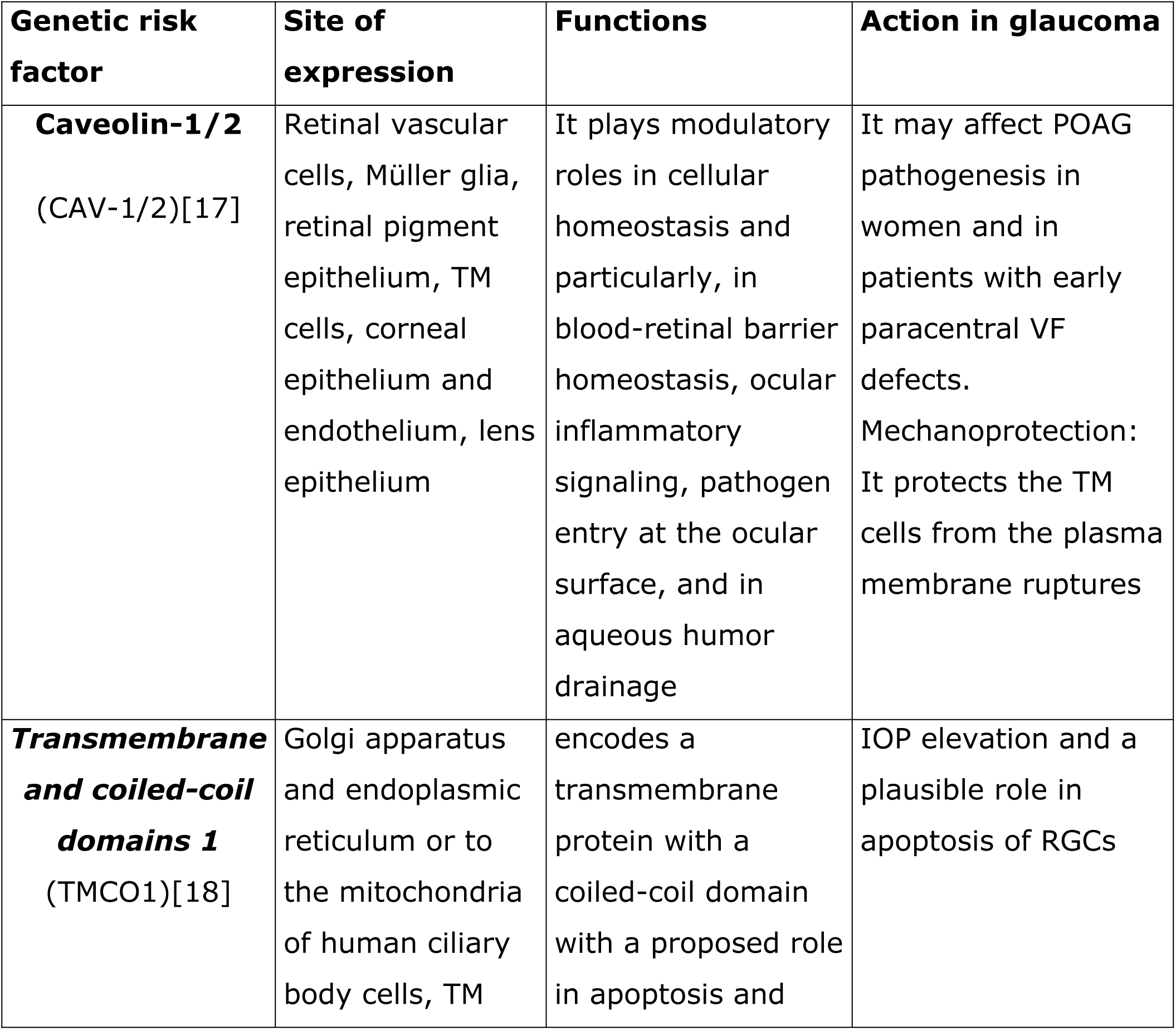
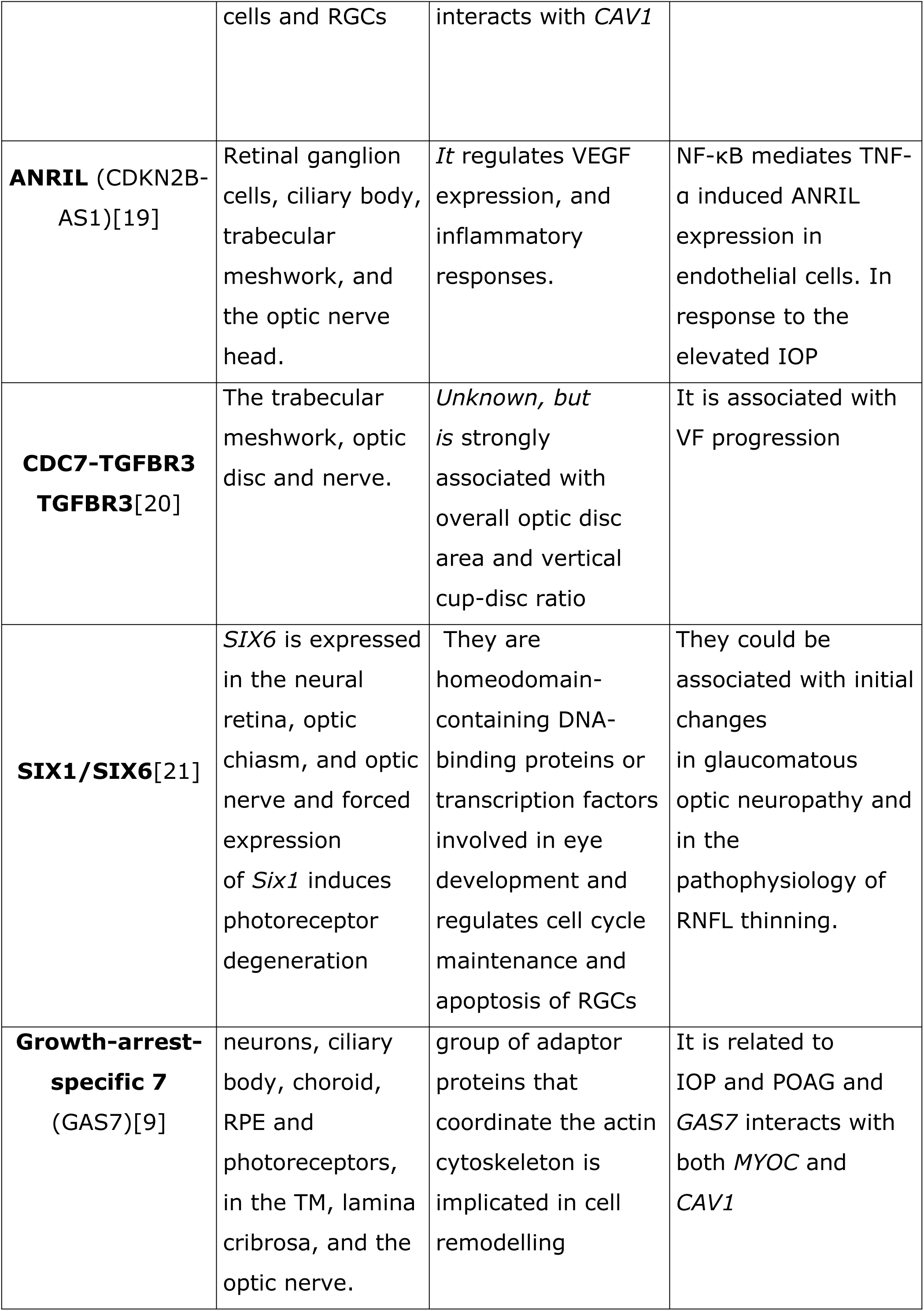
**Multifactorial Genetic Risk Factors:** in addition to well-known MYOCs and OPTNs, there are others reported below:

Individual susceptibility to the adverse effects of ROS is modulated by genetic polymorphisms of genes involved in their detoxification. This type of polymorphism is only one among the complex pathogenic events leading to glaucoma. Accordingly, adverse polymorphic variants of genes encoding for antioxidant activities only slightly influence the risk of developing the disease; the relative risk has been quantified as around 1.5 on comparing carriers of adverse genetic variants with carriers of wild-type variants, a situation referred to as ‘low penetrance’[14,15]. Nevertheless, these adverse genetic variants are highly frequent in humans, their prevalence being 35-40% in Caucasians. Evidence linking GSTM1 null polymorphisms to POAG exists in Asian but not in Caucasian populations, a situation probably due to the limited sample sizes so far examined[16]. For this reason, we conducted a survey on a sample of patients affected by glaucoma in Liguria (Italy) in order to ascertain the frequency of this polymorphism. The purpose of the present paper is to evaluate the impact of adverse genetic variants involved in defense against oxidative stress on the course and prognosis of glaucoma in a large patient cohort. Attention was focused on the gene encoding for GSTM1 because of the relevance of this function in neutralizing reactive oxygen species and the still open question of its role in glaucoma[13]. We examined the frequency of the GSTM1 null genotype in glaucoma patients in comparison with controls. We also examined the influence of this adverse polymorphism on the course of glaucoma.

## Methods

### 2.1 Study Groups

The study population comprised three group of patients: non-progressing POAGs well controlled with IOP-lowering medications (“Medical POAGs”); POAGs requiring surgery (“Surgical POAGs”); controls. Controls included age-matched non-glaucomatous healthy patients attending the outpatient clinic at our Institution and previously collected eye-bank specimens of non-glaucomatous patients as described in more detail elsewhere [3]. Only one eye per patients was considered for the analysis.

All included patients provided written informed consent. The study followed the tenets of Declaration of Helsinki and received IRB approval. The study was approved by the Ethics Committee of the AOU San Martino Genoa (Italy) on 20/01/2012, number 17/2011.

### 2.2 Primary Open Angle Glaucoma Diagnosis

The diagnosis of POAG based on slit-lamp examination, fundoscopy, standard automated perimetry (SAP) and, gonioscopy. Eyes were classified as glaucomatous if they had two consecutive reliable and repetable abnormal SAP test results in the presence of abnormal-appearing optic discs indicative of glaucoma by 90D lens fundus examination. Secondary glaucoma and angle closure glaucoma were excluded from the study. Ocular co-morbidity, systemic, or neurological disease and all other types of glaucoma represented exclusion criteria.

Medical POAGs comprised non-progressing patients defined by mean deviation change up to −0.5dB/year in the last three years with a minimum of five SAP and a controlled IOP. Surgical POAGs included patients undergoing surgery due to uncontrolled IOP on maximum tolerated medical therapy and unquestionable worsening of the VFs confirmed by al least two consecutive examinations.

### 2.3 Surgical procedure

All surgical patients underwent a standard trabeculectomy according to Cairns [22]. Careful dissection of the scleral flap was performed under the operating microscope to verify the presence of a pigmented corneal-scleral band in the trabecular meshwork before collecting it for analysis.

### 2.4 DNA extraction

DNA was extracted from oral swabs in medical POAGs and healthy controls while it was obtained directly from the trabecular meshwork of surgical POAGs and eye bank specimens as described in more detail elsewhere [3,23].

Tissue fragments were homogenized in the Tissuelyser (Qiagen) for 2 min at 30 Hz. DNA was purified using proteinase k digestion and affinity columns, using a commercially available kit (GenElute, Sigma, MO, USA).

The GSTM1 polymorphic deletion genotype was determined as described by Zhong et al. (Figure 1) [24]. The polymerase chain reaction (PCR) primers were P1 (59-CGC.CAT.CTT- .GTG.CTA.CAT.TGC.CCG-39), P2 (59-ATC.TTC.TCC.TCT-.TCT.GTC.TC-39), and P3 (59-TTC.TGG.ATT.GTA.GCA- .GAT.CA-39). P1 and P3 amplify a 230 bp product specific for the GSTM1 gene. Moreover, P1 and P2 amplify a 157 bp product specific for the GSTM4 gene, which, never having been deleted, was used as an internal control. PCR reaction was carried out in a total volume of 200 ml containing DNA, ATP, GTP, TTP, and CTP, DMSO, 50 mM MgCl2, and 2 U of PlatinumR Taq DNA Polymerase (Life Technologies, Rockville, Md.) in PCR buffer. The reaction was subjected to 35 cycles of amplification at 94°C for 1 min, 52°C for 1 min, and 72°C for 1 min by using a rotating thermal cycler (Rotorgene, Corbett ReseaRCH, Mortlake, Australia). The presence of the specific annealed hybrid (231 base pairs) was detected by SYBRGREEN staining. Melting curves were obtained by progressively increasing the temperature of the mixture from 70 to 98°C. The GSTM1 hybridized product has a melting temperature of 91.8°C. Accordingly, the presence of a peak at this temperature in the melting curve indicates the presence of a wild-type GSTM1 polymorphism, while the absence of a peak indicates the presence of a homozygous deletion for the GSTM1 gene.

**Figure 1:**
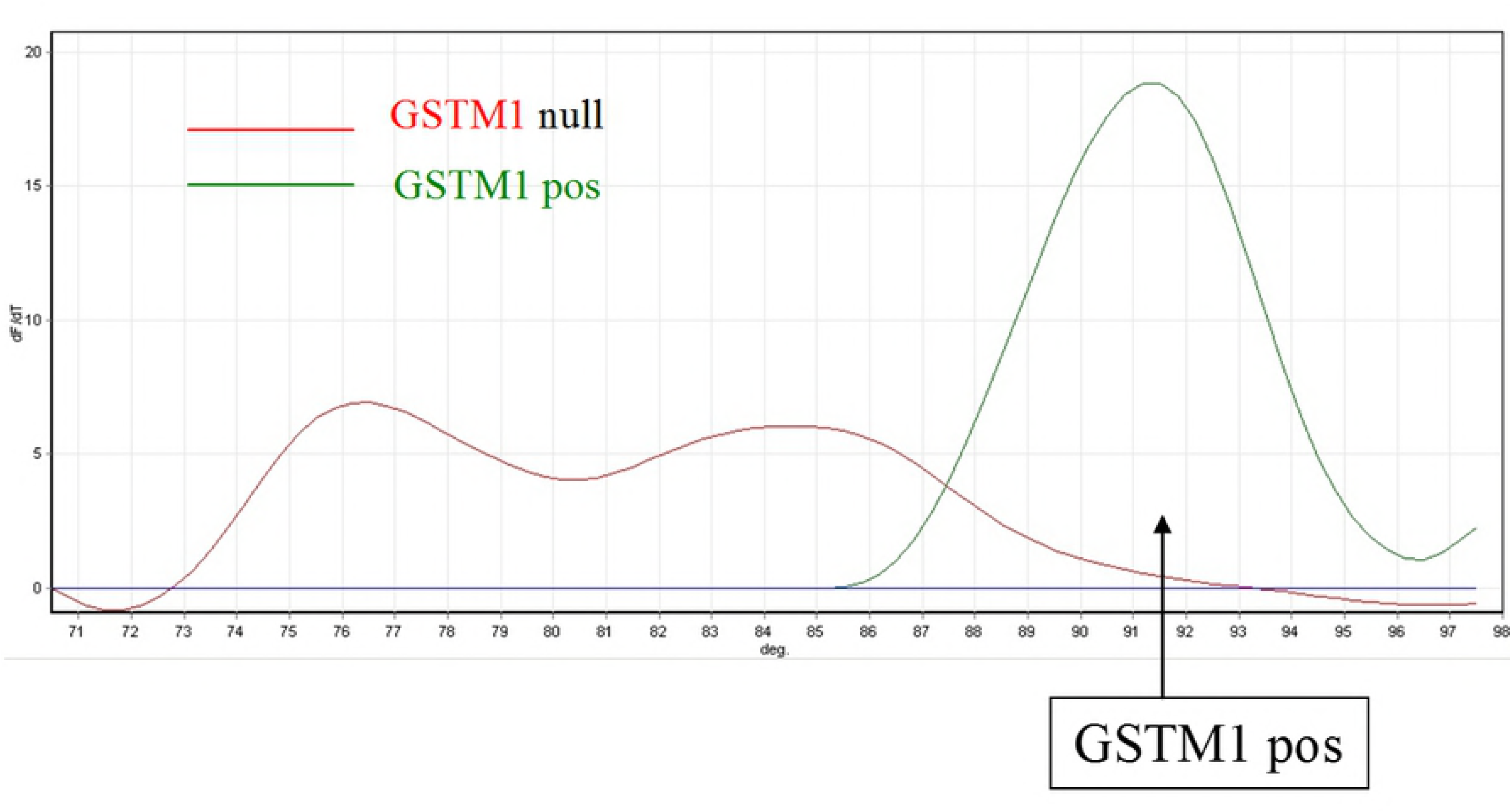
Detection of GSTM1 polymorphism by DNA melting curve analysis. GSTM1 gene sequence is amplified by PCR and double-strand annealed sequences labeled by fluorescent tracer (intensity of fluorescence signal, vertical axis). Temperature is progressively increased from 70 to 98°C (horizontal axis). The melting curve of amplified double-strand DNA (peaks) changes according to the sequence of nucleotides in the GSTM1 gene. GSTM1 positive polymorphism (green curve) corresponds to a melting temperature of 91.8°C; it cannot be detected in GSTM1 deleted polymorphism (red curve).

### 2.5 Statistical Analysis

In descriptive statistic, one-way analysis of variance was used for continuous data and Pearson’s chi-squared test when comparing frequencies among groups. Logistic regression assessed the associations of GSTM1 null genotype and medical or surgical POAGs with controls as the reference group. Odds ratios (ORs) and 95% confidence intervals (CIs) were also adjusted for age and gender. The Bonferroni correction was applied to correct for multiple testing. Statistical significance was set at alpha = 5%. Based on the work of Lavaris et al., with 104 cases and 198 controls, the study has a power of 80% and is sufficiently powered[25]. All statistical analyses were performed using Stata 15 (Statacorp LCC, College Station, TX).

## Results

GSTM1 polymorphism detection was achieved in all samples. Table 2 reports the demographics of POAGs and controls: no significant difference regarding gender frequencies and mean age were detected among groups (p=0.950 and p=0.275, respectively). All the enrolled subjects were Caucasian. Table 3 reports the of the GSTM1 genotype frequencies for each group. A chi-square test shows that a significative relationship exists between GSTM1 status and groups (p=0.033). Figure 2 shows the percentage of the GSTM1 null status in the three groups. Logistic regression using the disease status as the binary outcome and the GSTM1 genotype as the predictor variable show that a positive association exist for surgical POAGs but not for medical POAG (p=0.028 and p=0.20, respectively). Table 4 shows the odds ratio adjusted for age and gender.

**Table 2:** Demographics

|  | Medical POAGs<br>(n=104) | Surgical POAGs<br>(n=158) | Controls<br>(n=263) |  |
| --- | --- | --- | --- | --- |
| Gender, % (n) |  |  |  |  |
| Female | 54 (51.9) | 83 (52.5) | 134 (51.0) | 0.950<sup>a</sup> |
| Male | 50 (48.1) | 75 (47.5) | 129 (49.0) |  |
| Age, years | 68.3 (7.1) | 69.8 (3.9) | 69.1 (6.5) | 0.275<sup>b</sup> |

Data are number (%) or mean (SD). POAG: primary open angle glaucoma. <sup>a</sup> Chi-squared test. <sup>b</sup> One-way analysis of variance.

**Table 3:** GSTM1 genotype among POAGs and Controls

| GSTM1 status | Controls | Medical<br>POAGs | Surgical<br>POAGs |
| --- | --- | --- | --- |
| positive | 144 (54.75) | 47 (45.19) | 67 (42.41) |
| null | 119 (45.25) | 57 (54.81) | 91 (57.59) |
| Total | 263 | 104 | 158 |

Data are number (%). GSTM1: glutathione transferase isoenzyme M1; POAG: Primary open angle glaucoma. Chi-squared test, p=0.033

**Table 4:** Odds ratio adjusted for gender and age

| Gene<br>Status | No. of<br>exposed<br>Controls | Medical POAGs |  |  | Surgical POAGs |  |  |
| --- | --- | --- | --- | --- | --- | --- | --- |
|  |  | No. of<br>exposed<br>cases | OR (95%<br>CI) | p | No. of<br>exposed<br>cases | OR (95%<br>CI) | p |
| GSTM1<br>positive |  |  | reference |  |  | reference |  |
| GSTM1<br>null | 119 | 57 | 1.44<br>(0.86 to<br>2.39) | 0.326* | 91 | 2.01<br>(1.26 to<br>3.21) | 0.006* |

GSTM1: glutathione transferase isoenzyme M1; POAG: Primary open angle glaucoma; OR: Odds ratio. \* with multiple testing correction.

**Figure 2:**
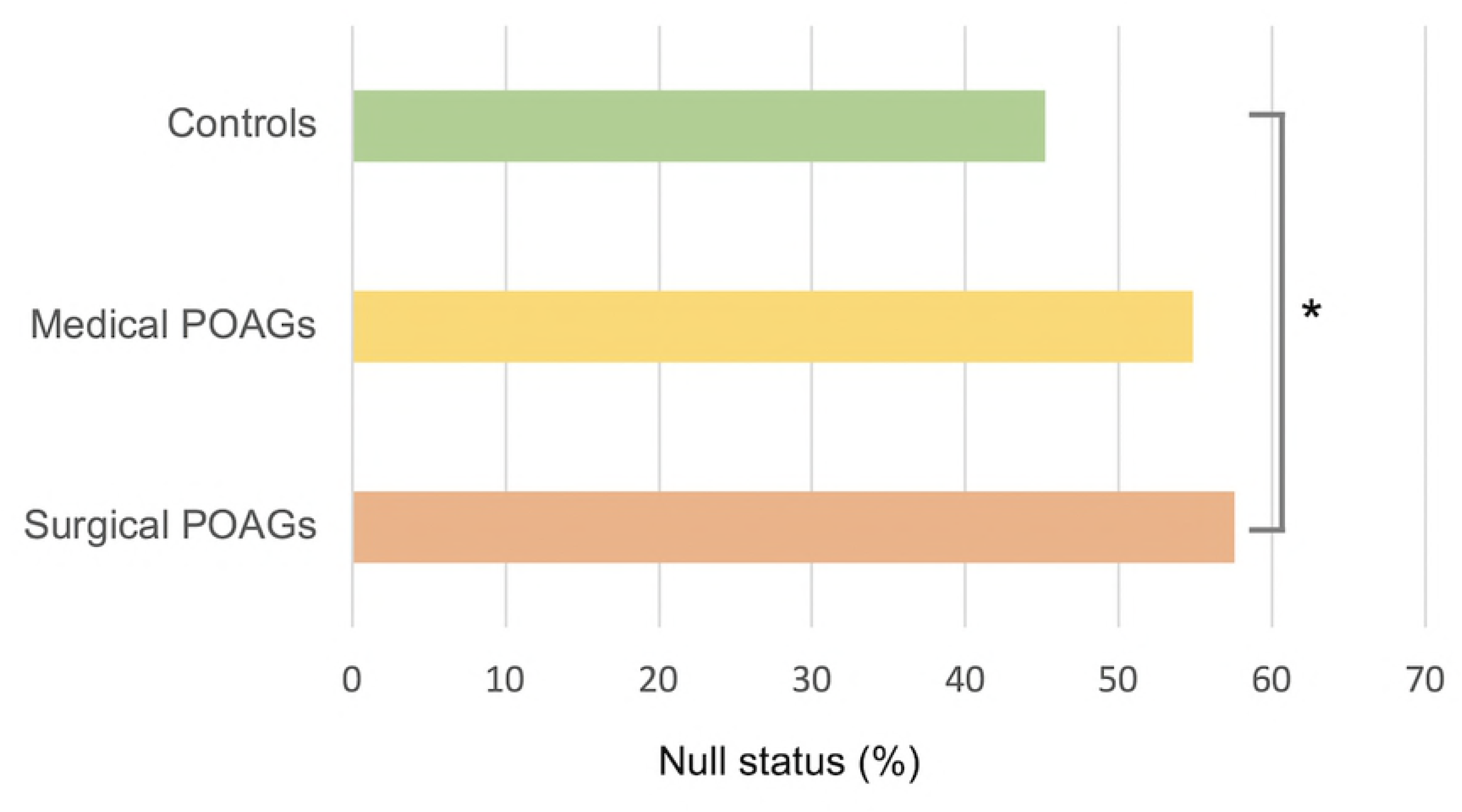
GSTM1 null status among the three groups. * statistically significant

## Discussion

Glaucoma is a neurodegenerative disease and many factors, including free radicals, are involved in its pathogenesis[8]. One of the main sources of free radicals is ultraviolet rays, which are able to alter nucleic acids, membranes and cellular functions[26]. They can also activate pathways that lead to inflammation, such as the NFkB pathway. Furthermore, this pathway is able to induce the transcription of a myriad of genes that mediate diverse cellular processes, such as immunity, inflammation, proliferation, apoptosis and cellular senescence[27]. Ultraviolet light does not directly reach the angle of the anterior chamber. However, ROS are able to affect the cellularity of the human TM[28], and the conventional aqueous outflow is more susceptible to oxidative damage than other tissues of the anterior chamber[29]. Oxidative damage, as measured directly on the TM, is much greater in glaucomatous subjects than in controls and is directly proportional to IOP levels and visual-field defects[8,29]. Furthermore, oxidative DNA damage in the TM is significantly correlated with age showing striking morphological decay. In addition, the genotype of senile trabecular cells is markedly increased[30]. The inhibition of contractility of the TM cells occurs with age[31], and there is a marked loss of TM cells[32]. An important source of free radicals is the mitochondria, in which oxidative damage is the result of an error in mitochondrial DNA replication. This mitochondrial deletion causes an energy deficit and cellular atrophy[3]. In the conventional outflow pathway, the mitochondrial deletion that occurs during glaucoma is much greater than in healthy patients. Mitochondrial deletion occurs only in POAG and in pseudoexfoliative glaucoma; it is not found in other types of glaucoma, in which the pathogenesis is different[7]. Thus, oxidative stress causes alterations of DNA and RNA; as a result, there will be changes in proteins and microRNA. MicroRNAs are recognized as important post-transcriptional regulators of gene expression, and the difference in microRNA between the glaucomatous aqueous humor and that of controls is very evident[33]. Even the proteins found in the aqueous humor of glaucoma patients are different from those of healthy subjects. The expression of these proteins in the aqueous humor of glaucoma patients reflects the damage occurring in the endothelia of the anterior chamber, particularly to their cytoskeleton. The endothelium of the TM acts like that of small vessels or capillaries[34]. Thus, during open-angle glaucoma, chronic damage to the TM occurs, which is reflected in the protein expression of trabecular cells that flow in the aqueous humor. These accurately describe the cascade of events that starts with oxidative damage and first leads to the malfunction of the TM and then to IOP increase[6]. These proteins can become biological signals that reach the posterior segment. Indeed, in 1986, Smith discovered the pathway that leads from the anterior chamber to the optic nerve head[35]. So, the Nestin, is a protein probably expressed in the anterior chamber in response to the TM malfunction, and serves to activate the stem cells of the TM; however, when it reaches the posterior segment, it actives the glia. As AKAP 2 in the anterior chamber, reflects the damage to TM motility, while in the posterior segment, it may be an intracellular signal that triggers RGC death by apoptosis[31]. In this scenario, the lack of the gene GSTM1 could have a pathogenetic role. Indeed, individuals with GSTM1 deficiency show a greater level of DNA damage[36]. The differential distribution of polymorphic gene variants in different human populations around the world may influence the environmental diseases which they acquire[37]. The frequency of the GSTM1 null genotype in humans ranges from 30-50%, depending on the ethnic origin of the individual[38]. The variations in GSTM1 expression among control populations makes it very difficult for researchers to select suitable control groups to be used in order to determine the association between GST genotype and disease susceptibility. With regard to glaucoma, its incidence varies according to the population studied. Thus, the null GSTM1 genotype has been reported to be more common in POAG patients in both a Turkish and an Italian population[39]. A meta-analysis conducted by Huang et al. suggested that GSTM1 null genotypes were associated with an increased POAG risk in Asian populations, but not in Caucasian or mixed populations, although the studies they analyzed had limited sample sizes[16]. A further meta-analysis indicated that *GST* polymorphisms may contribute to the increased risk of glaucoma, and still another, conducted in the same year, reported that there might be a significant association between the GSTM1 null genotype and POAG risk in East Asians[40]. The disparity of these studies may certainly be due to genetic variations related to race. Nevertheless, we think that it should be evaluated whether or not this polymorphism affects not only the overall frequency of glaucoma, but also the trend in the multiple clinical variables that characterize the prognosis of glaucoma patients. For this reason, we evaluated the frequency of the GSTM1 null genotype in two different clinical groups of Caucasian patients: glaucoma tonometrically compensated by pharmacological therapy, and more severe uncompensated glaucoma subjected to filtration surgery. Our results provide evidence that the homozygous deletion is associated with severe POAG, i.e. unresponsive to medical treatments and requiring surgical therapy. Accordingly, GSTM1 deletion may constitute an adverse risk factor for POAG prognosis that can be evaluated by means of non-invasive sampling (oral swab).

The GSTM1 null genotype is a risk factor because this genotype causes an increase in oxidative damage. Indeed, reduced GST function might interfere with the metabolism of oxidative intermediates and exacerbate the damaging effects of oxidative stress on the optic nerve, thus contributing to glaucomatous neurodegeneration[41]. Rocha et al. found that in the Brazilian population the GSTM1 null genotype was associated with higher IOP levels and more severe damage to the optic nerve and visual field[42]. Indeed, extensive and repeated oxidative stress *in vivo* can reduce TM cell adhesion, resulting in cell loss and compromising TM integrity[43].[27][30] As the loss of trabecular function is of fundamental importance in the induction of optic nerve damage[31] it is therefore reasonable that the GSTM1 null genotype induces worsening of the clinical development of glaucoma.

Our research shows that, in the Caucasian population, the GSTM1 null genotype is a risk factor in those affected by glaucoma, is more often observed in patients who require surgery. In conclusion, according to our research, the GSTM1null genotype in glaucoma patients is an indication for the use of both topical and systemic antioxidants to protect both the trabecular meshwork and the optic nerve head from oxidative injury. Further studies will be performed in order to evaluate the impact of this type of therapy in these patients.

